# Leveraging genome-wide data to investigate differences between opioid use *vs*. opioid dependence in 41,176 individuals from the Psychiatric Genomics Consortium

**DOI:** 10.1101/765065

**Authors:** Renato Polimanti, Raymond K. Walters, Emma C. Johnson, Jeanette N. McClintick, Amy E. Adkins, Daniel E. Adkins, Silviu-Alin Bacanu, Laura J. Bierut, Tim B. Bigdeli, Sandra Brown, Kathy Bucholz, William E. Copeland, E. Jane Costello, Louisa Degenhardt, Lindsay A Farrer, Tatiana M. Foroud, Louis Fox, Alison M. Goate, Richard Grucza, Laura M. Hack, Dana B. Hancock, Sarah M. Hartz, Andrew C. Heath, John K. Hewitt, Christian J. Hopfer, Eric O. Johnson, Kenneth S. Kendler, Henry R. Kranzler, Ken Krauter, Dongbing Lai, Pamela A. F. Madden, Nicholas G. Martin, Hermine H. Maes, Elliot C. Nelson, Roseann E. Peterson, Bernice Porjesz, Brien P. Riley, Nancy Saccone, Michael Stallings, Tamara Wall, Bradley T. Webb, Leah Wetherill the Psychiatric Genomics Consortium Substance Use Disorders Workgroup, Howard J. Edenberg, Arpana Agrawal, Joel Gelernter

## Abstract

To provide novel insights into the biology of opioid dependence (OD) and opioid use (i.e., exposure, OE), we completed a genome-wide analysis comparing up to 4,503 OD cases, 4,173 opioid-exposed controls, and 32,500 opioid-unexposed controls. Among the variants identified, rs9291211 was associated with OE (a comparison of exposed vs. unexposed controls; z=-5.39, p=7.2×10^−8^). This variant regulates the transcriptomic profiles of *SLC30A9* and *BEND4* in multiple brain tissues and was previously associated with depression, alcohol consumption, and neuroticism. A phenome-wide scan of rs9291211 in the UK Biobank (N>360,000) found association of this variant with propensity to use dietary supplements (p=1.68×10^−8^). With respect to the same OE phenotype in the gene-based analysis, we identified *SDCCAG8* (z=4.69, p=10^−6^), which was previously associated with educational attainment, risk-taking behaviors, and schizophrenia. In addition, rs201123820 showed a genome-wide significant difference between OD cases and unexposed controls (z=5.55, p=2.9×10^−8^) and a significant association with musculoskeletal disorders in the UK Biobank (p=4.88×10^−7^). A polygenic risk score (PRS) based on a GWAS of risk-tolerance (N=466,571) was positively associated with OD (OD cases vs. unexposed controls, p=8.1×10^−5^; OD cases vs. exposed controls, p=0.054) and OE (exposed controls vs. unexposed controls, p=3.6×10^−5^). A PRS based on a GWAS of neuroticism (N=390,278) was positively associated with OD (OD cases vs. unexposed controls, p=3.2×10^−5^; OD cases vs. exposed controls, p=0.002) but not with OE (p=0.671). Our analyses highlight the difference between dependence and exposure and the importance of considering the definition of controls (exposed vs. unexposed) in studies of addiction.

## INTRODUCTION

The prevalence of opioid dependence (OD) is at epidemic levels and significantly affects public health and social and economic well-being. The use of opioid medications for analgesia is common, and opioids are considered a gold standard for pain control. However, they are also highly addictive, and are, along with heroin^1^, the leading contributors to the ongoing epidemic of opioid misuse and the high rate of fatal overdoses from prescription opioids^2–4^.

Understanding the biology of human responses to opioids may lead to effective preventive strategies and treatments to reduce OD and its harmful consequences. Human genetic research has the potential to dissect the basis of inter-individual variability in the response to opioid exposure (i.e., whether an individual develops dependence on opioids). Genome-wide association studies (GWAS) of large cohorts have identified a number of risk loci and molecular pathways involved in the predisposition to numerous psychiatric disorders and behavioral traits^5, 6^. Previous OD GWAS included up to 10,000 participants and identified genome-wide significant (GWS) associations in *KCNG2*, *KCNC1*, *APBB2*, *CNIH3*, and *RGMA*^7–10^. However, there was no consistency across the previous OD GWAS with respect to the individual GWS loci, probably due to the limited statistical power and differences in case and control definitions in the context of polygenic architecture (thousand causal loci with small effect).

A key potential contributor to the lack of consistency in findings from prior GWAS is that different study designs were used. The most relevant design variation is related to the assessment of opioid exposure in controls where two different control definitions have been considered: i) individuals exposed to opioids (OE) medically or illegally who did not develop OD; or ii) individuals without an OD diagnosis who were not assessed for opioid exposure. Although including individuals not exposed to opioids in the control group increases the overall sample size, it also potentially adds noise by including individuals who would have been likely to become OD if exposed, given the highly addictive nature of opioid drugs. Furthermore, exposure to opioids is a behavioral trait *per se*, and likely to be associated with its own specific genetic architecture, which may be different between licit and illicit exposure. Opioid use is rarer than the use of many other substances and it is often observed in individuals affected by severe mental and physical illnesses^11, 12^. Comparisons of OD cases with predominantly unexposed controls is more likely to confound genetic risk for exposure to opioids with genetic factors specific to the transition to OD. Indeed, at least one prior smaller GWAS^7^ found that comparisons of OD cases to controls with significant exposure and from similar neighborhoods resulted in a GWS finding while comparisons with general population controls did not identify any GWS variants.

We leveraged genotypic and phenotypic information from 41,176 participants from 11 studies that are part of the Psychiatric Genomics Consortium (PGC) SUD working group to investigate genetic differences between OD cases (n=4,503), OE controls (n=4,173), and opioid-unexposed (OU) controls (n=32,500) using GWAS and polygenic risk score (PRS) analyses. In addition to identifying loci related to OD and OE, we also examined whether OD and OE could be differentiated with respect to their relationship with genetic liability to risk-taking behaviors and negative personality features (i.e., neuroticism), to provide further insights into the genetic architecture underlying opioid use and misuse.

## MATERIALS AND METHODS

### Cohorts

Of the 11 studies from the PGC-SUD workgroup, seven were case-control studies and four were family-based studies (Supplementary Table 1; Supplementary Methods). Lifetime OD diagnoses was based on DSM-IV OD criteria^13^ and were derived either from clinician ratings or semi-structured interviews. The two control groups included OE controls (individuals without a lifetime OD diagnosis who were exposed to opioids at least once) and OU controls (individuals with no lifetime OD diagnosis who were not exposed to opioids). Lifetime opioid exposure included both licit, prescribed opioids and those used outside appropriate medical care. Some, but not all, studies distinguished between these forms of exposure. This study, which involved the analysis of de-identified data, was approved by the institutional review board (IRB) at Yale University School of Medicine and was conducted in accordance with all relevant ethical regulations. Each contributing study obtained informed consent from participants and ethics approvals for their study protocols from their respective review boards in accordance with applicable regulations.

### Quality control and imputation

Individual genotype information was available for each subject. The Ricopili pipeline^14^ (https://github.com/Nealelab/ricopili) was used for the QC and imputation of the case-control cohorts. Most family-based cohorts were analyzed with the Picopili pipeline^15^ (https://github.com/Nealelab/picopili), which is designed to conduct genome-wide meta-analyses accounting for family structure. The genetic data from the Collaborative Studies on the Genetics of Alcoholism were imputed independently as previously described^16^ because of the need in that study to merge data on members of large multiplex families who were genotyped across multiple genotyping arrays.

Details regarding the QC criteria were reported previously^15^. Briefly, after initial sample and variant QC, population outlier samples were excluded, and each retained individual was assigned to a specific ancestry on the basis of the principal components derived from genome-wide data. The 1000 Genomes Project Phase 3 reference panel^17^ was used as a reference for the ancestry assignment. Based on genetic information, we identified 9,591 and 31,585 individuals of African and European descent, respectively. Other ancestry groups were not investigated due to the limited number of informative subjects. The final QC criteria included variant filters for call rate, heterozygosity, and departure from Hardy–Weinberg equilibrium expectations (HWE), performed within each ancestry group in each cohort stratified by genotyping array. We also used sample QC filters for cryptic relatedness and for departures from reported pedigree structures.

Imputation was performed using SHAPEIT2^18^ and IMPUTE2^19^, and the 1000 Genomes Project Phase 3 reference panel, which includes five continental groups^17^. High-quality imputed SNPs were retained for the association analysis, filtering for imputation INFO score > 0.8 and minor allele frequency (MAF) > 0.01 before analysis. After imputation, we tested for duplicated samples and cryptic relatedness among the cohorts analyzed. The association analysis was conducted considering variants present in at least 80% of the cohorts investigated (Supplementary Table 2).

### Data analysis

The association analysis was conducted stratifying each cohort by ancestry (i.e., African and European ancestries) and genotyping array. For case–control studies, imputed dosages were entered in a logistic regression. For family-based studies, logistic mixed models were used. The association analyses were adjusted for sex and the within-ancestry top 10 principal components to account for possible confounding by population stratification. To investigate differences between OE and OD, three phenotype definitions were considered: i) OD cases vs. OE controls (OD_exposed_; n=4,503 and 4,173, respectively); ii) OD cases vs. OU controls (OD_unexposed_; n=4,238 and 17,700, respectively; as explained in the Supplementary Methods, the reduction of sample size is due to the removal of cohorts with low case-control ratio); OE controls vs. OU controls (OE_controls_; n=4,173 and 32,500). For each phenotype, meta-analyses of the results across the different cohorts were conducted in METAL with weights proportional to the square-root of the sample size for each study^20^. The effective sample size of each cohort was calculated on the case-control ration and the relatedness matrix. Ancestry-specific (African-ancestry and European-ancestry) and trans-ancestry meta-analyses were conducted. Heterogeneity was evaluated across all cohorts and between study designs.

To investigate the loci identified in the individual GWAS further, we performed a phenome-wide scan considering 4,082 traits assessed in up to 361,194 participants from the UK Biobank using previously generated GWAS association summary data^21^. Details regarding QC criteria and GWAS methods of this previous analysis are available at https://github.com/Nealelab/UK_Biobank_GWAS/tree/master/imputed-v2-gwas. Briefly, the association analyses for all phenotypes were conducted using regression models available in Hail (available at https://github.com/hail-is/hail) including the first 20 ancestry principal components, sex, age, age^2^, sex×age, and sex×age^2^ as covariates. We applied a false discovery rate (FDR) multiple testing correction (q<0.05) to account for the number of variants and phenotypes tested.

Linkage disequilibrium (LD) score regression^22^ was performed to estimate the heritability explained by common SNPs (h^2^_g_) in the European-ancestry meta-analysis of case-control and family-based cohorts. The inclusion of related subjects may affect the LD score regression results due to the residual effect of family structure on the summary association data. To limit this potential confounder, the analyses was limited to variants assessed in more than 80% of the total sample and considering the effective sample size adjusted for both case-control ratio and family structure. The heritability analysis was not conducted on African-specific and trans-ancestry meta-analyses, because LD score regression is not suitable when analyzing GWAS summary data derived from admixed populations^22^. LD score regression analysis was performed considering HapMap3 SNPs^23^ and LD scores computed from the 1000 Genomes Project reference for European populations. Conversion of h^2^_g_ estimates from observed scale to liability scale was performed accounting for the difference between population prevalence (OD_exposed_=1%, OD_unexposed_=1%, and OE_controls_=5%) and sample prevalence (OD_exposed_=55%, OD_unexposed_=22%, and OE_controls_=12%).

Gene-based association, enrichment analysis for molecular pathways, Gene Ontologies (GO) annotations and tissue-specific transcriptomic profiles were conducted using the MAGMA tool^24^ implemented in the FUMA platform^25^. Information regarding molecular pathways and GO annotations was derived from MsigDB v6.2^26^. Tissue-specific transcriptomic profiles were derived from GTEx V7^27^ and BrainSpan^28^. A Bonferroni multiple testing correction was used to control for the number of tests conducted in each enrichment analysis. GTEx data were also used to verify whether the GWS loci identified affect the transcriptomic regulation of the surrounding genes. To evaluate the effect across multiple tissues, we considered multi-tissue expression quantitative trait locus (eQTL) data. These were calculated using Meta-Tissue ^29^. This meta-analytic approach calculates a posterior probability (m value) that an effect exists in each of the tissues tested assuming that the eQTL effect is consistent across the affected tissues. M values>0.9 indicate that the tissue was predicted to show the eQTL association.

A PRS analysis was conducted to test the genetic overlap with behavioral traits that could differentiate between OD and OE status using the PRSice software^30^. Risk-taking and neuroticism were selected as we expected they would capture genetic susceptibility to early versus later stages of opioid use and misuse. For polygenic profile scoring, we used summary statistics generated from large-scale GWAS of risk tolerance (N= 466,571)^31^ and neuroticism (N=390,278)^32^. We considered multiple association P value thresholds (P_T_ < 5×10^−8^, 10^−7^, 10^−6^, 10^−5^, 10^−4^, 0.001, 0.05, 0.1, 0.3, 0.5, 1) for SNP inclusion. The PRS were calculated after using P-value-informed clumping with a LD cut-off of R^2^ = 0.3 within a 500 kb window and excluding the major histocompatibility complex region of the genome because of its complex LD structure. The PRS were calculated considering unrelated subjects of European descent available in both case-control and family-based cohorts (OD_exposed_ N_effective_=3,038; OD_unexposed_ N_effective_=4,728; and OE_controls_ N_effective_=5,376). The PRS were fitted in regression models with adjustments for sex and the top 10 within-ancestry principal components. We applied FDR multiple testing correction (q<0.05) to correct for the number of thresholds tested.

## RESULTS

### GWAS Analyses Comparing OD and OE Traits

The GWAS meta-analyses of OD_exposed_, OD_unexposed_, and OE_controls_ phenotypes included up to 4,503 OD cases (African-ancestry=1,231; European-ancestry=3,272), 4,173 OE controls (African-ancestry=1,297; European-ancestry=2,876), and 32,500 OU controls (African-ancestry=7,063; European-ancestry=25,437). Significant SNP-heritability was observed for OD_unexposed_ (liability-scale h^2^_g_=0.28, SE=0.1; population prevalence=0.01, sample prevalence=0.22), but not for OD_exposed_ (liability-scale h^2^_g_=-0.08, SE=0.08; population prevalence=0.01, sample prevalence=0.55) and OE_controls_ (liability-scale h^2^_g_=0.05, SE=0.1; population prevalence=0.05, sample prevalence=0.12). Moderate genome-wide inflation was observed in the European-specific meta-analyses, but genomic control using lambda or the LD score regression intercept did not affect the significance of any variants in the GWAS meta-analyses (Supplementary Table 3; Supplementary Methods).

In the OD_exposed_ analysis, which is the most relevant comparison to dependence liability given exposure but that most constricted the sample size, no association survived the genome-wide significance threshold (P=5×10^−8^). Additionally, there were no significant enrichments for GO annotations, molecular pathways, nor tissue-specific regulation.

The other two comparisons included the much larger OU group. The OD_unexposed_ comparison identified a GWS association in the African-ancestry meta-analysis, rs201123820 on chromosome 18 (z=5.55, p=2.9×10^−8^; Figure 1A; Table 2; see Supplementary Table 4 for ancestry-specific results for each genome-wide significant variant). With respect to this locus, no heterogeneity was observed among the cohorts included in the meta-analysis (heterogeneity: I^2^=0, p=0.473; Supplementary Table 5). The gene-based association analysis identified a GWS gene in the same genomic region, *C18orf32* (p=1.8×10^−6^; Supplementary Figure 1A). Additionally, in the African-ancestry meta-analysis, we also observed an enrichment for adipose tissue (beta=0.04, p=4.21×10^−4^; Figure 2A) and GO:0034498 – *early endosome to Golgi transport* (beta=1.01, p=5.1×10^−8^). In the trans-ancestry meta-analysis, we observed significant enrichment for specific adult stages of brain development (37 y; beta=0.06, p=6.22×10^− 4^; 15 y; beta=0.06, p=0.001; 36yrs: beta=0.06, p=0.002; Figure 2B) and GO:0007143∼*female meiotic division* (beta=0.73, p=1.08×10^−7^). In the European-ancestry OD_unexposed_ GWAS meta-analysis, no result survived multiple testing correction. Furthermore, variants identified in prior opioid dependence GWAS were not genome-wide significant in this study (Supplementary Table 6).

**Table 1:** Top loci identified considering different OD and OE phenotypic definitions.

| Single-variant Associations |  |  |  |  |  |  |  |  |  |
| --- | --- | --- | --- | --- | --- | --- | --- | --- | --- |
| Phenotype | Meta-analysis | rsid | Chromosome | Location (bp) | Effect Allele | Other Allele | Effect Allele Frequency | Z score | P value |
| OD<sub>unexposed</sub> | AFR | rs201123820 | 18 | 47025347 | T | TAAACAAAAACA | 0.019 | 5.547 | 2.90E-08 |
| OE<sub>controls</sub> | EUR | rs9291211 | 4 | 42139132 | A | G | 0.782 | -5.387 | 7.16E-08 |
|  | Trans-ancestry | rs12461856 | 19 | 55433852 | A | G | 0.8402 | -5.606 | 2.07E-08 |
| Gene-based Associations |  |  |  |  |  |  |  |  |  |
| Phenotype | Meta-analysis | Gene | Chromosome | Location-Start (bp) | Location-End (bp) | SNP (n) | Z score | P value |  |
| OD<sub>unexposed</sub> | AFR | *C18orf32* | 18 | 47008028 | 47013622 | 18 | 4.633 | 2E-06 |  |
| OE<sub>controls</sub> | EUR | *BEND4* | 4 | 42112955 | 42154895 | 81 | 4.756 | 1E-06 |  |
|  | Trans-ancestry | *SDCCAG8* | 1 | 243419320 | 243663394 | 407 | 4.686 | 1E-06 |  |

**Figure 1:**
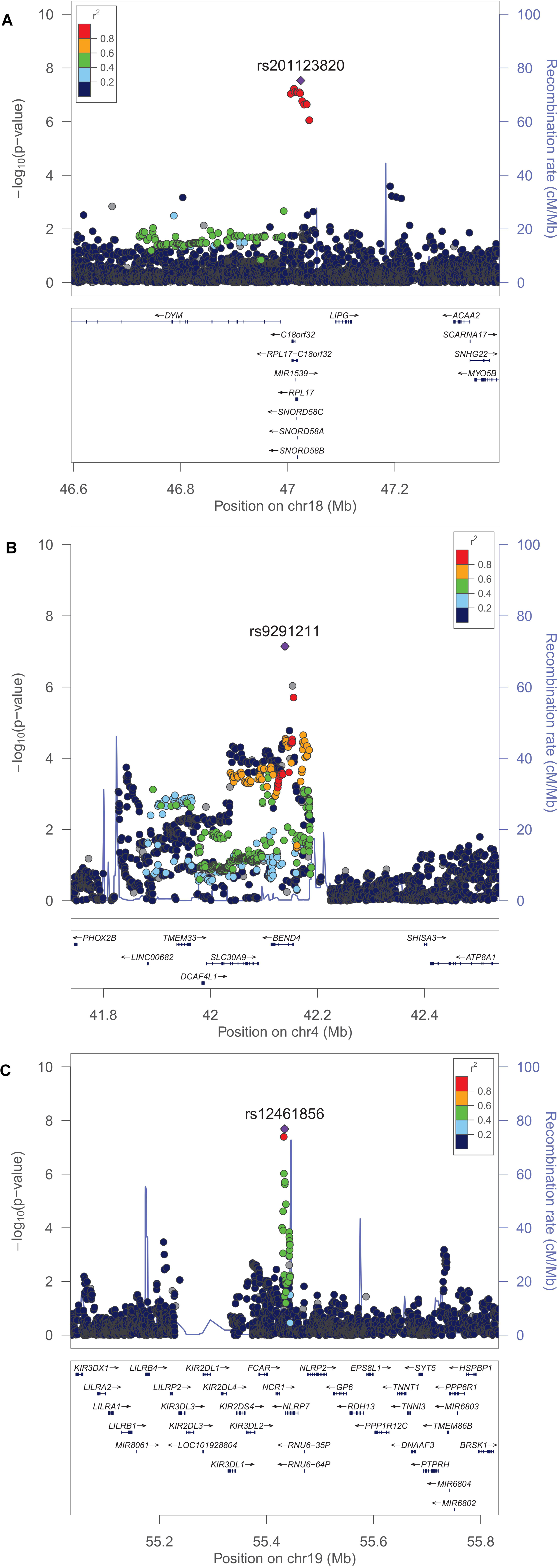
Regional Manhattan plots of the genetic association identified: rs201123820 in the African-ancestry OD_unexposed_ GWAS meta-analysis (**A**); rs92911211 in the European-ancestry OE_controls_ GWAS meta-analysis (**B**); rs12461856 in the trans-ancestry OE_controls_ GWAS meta-analysis (**C**).

**Figure 2:**
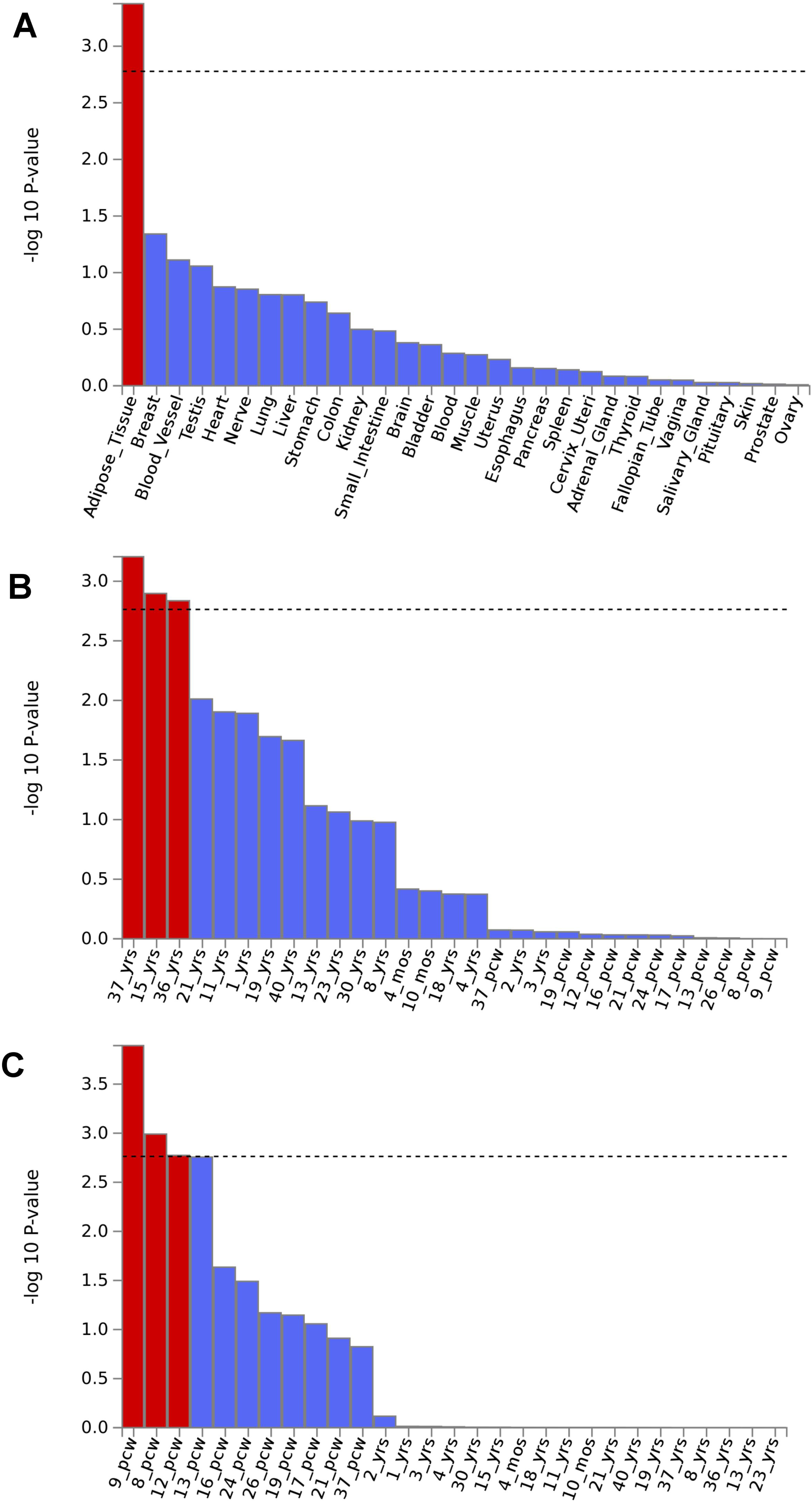
Significant tissue enrichments identified in the African-ancestry OD_unexposed_ meta-analysis (**A**); the trans-ancestry OD_unexposed_ GWAS meta-analysis (**B**), and the European-ancestry OE_controls_ GWAS meta-analysis (**C**).

In GWAS meta-analysis of OE_controls_ in the European-ancestry cohort, we identified a genetic association that reached “suggestive” significance (p<9×10^−8^), rs9291211 on chromosome 4 (z=-5.38, p=7.2×10^−8^; Figure 1B). With respect to this locus, no heterogeneity was observed among the cohorts included in the meta-analysis (heterogeneity: I^2^=0, p=0.879; Supplementary Table 5). This variant (or LD proxies in the same ancestry group) was identified in previous GWAS of behavioral traits: alcohol consumption (rs4501255, LD proxy r^2^=0.94, p=5×10^−10^)^33^; neuroticism (rs9291211, p=2×10^−8^)^32^; and helping behavior (rs2880666, LD-proxy r^2^=0.77, p=5×10^−7^)^34^. Additionally, rs9291211 is an eQTL for *SLC30A9* and *BEND4* in multiple tissues (GTEx multi-tissue eQTL p = 1.2×10^−26^ and 2.88×10^−9^, respectively). The rs9291211×*SLC30A9* eQTL (i.e., rs9291211 regulating the *SLC30A9* expression) showed a posterior probability>90% in seven brain tissues (amygdala m=0.99; anterior cingulate cortex m=1; caudate m=1; cortex m=0.99; hypothalamus m=1; nucleus accumbens m=1, putamen m=0.99; Supplementary Figure 2A). The rs9291211×*BEND4* eQTL showed posterior probabilities>90% in two brain tissues (caudate m=0.9; cortex m=1; Supplementary Figure 2B). *BEND4* was also identified as GWS in the gene-based test (p=9.9×10^−6^; Supplementary Figure 1B). The European-ancestry OE_controls_ meta-analysis also showed enrichment for several brain development stages: post-conception weeks 9 (beta=0.04, p=1.28×10^−4^), 8 (beta=0.032, p=0.001), and 12 (beta=0.04, p=0.002) (Figure 2C). No result in the association and the enrichment analyses based on the African-ancestry OE_controls_ GWAS meta-analysis survived multiple testing correction.

In the trans-ancestry GWAS meta-analysis of OE_controls_, we observed an additional single-variant GWS association, rs12461856 on chromosome 19 (z=-5.61, p=2.1×10^−8^; Figure 1C). With respect to this locus, no heterogeneity was observed among the cohorts included in the meta-analysis (heterogeneity: I^2^=0, p=0.554; Supplementary Table 5). A gene-based GWS association was identified for *SDCCAG8* on chromosome 1 (p=1.4×10^−6^, Supplementary Figure 1C). Significant enrichments were also observed for GO:0017069∼*small RNA binding* (beta=0.65, p=5.4×10^−7^) and a curated gene set related to genes downregulated 6 h after induction of *HoxA5* expression in a breast cancer cell line (Standard name: CHEN_HOXA5_TARGETS_6HR_DN; beta=1.37, p=8.6×10^−6^).

### Phenome-wide Scan in UK Biobank

To extend the phenotypic breadth of our findings, we conducted a phenome-wide scan (4,082 traits tested; Supplementary Table 7) for three variants identified in 361,194 participants of European descent from the UK Biobank. Rs201123820 was identified in the African-ancestry OD_unexposed_ GWAS meta-analysis. Although this variant was not significant in the European-ancestry meta-analysis (Supplementary Table 4), Rs201123820 has reasonably similar MAF in African and European populations (1000 Genomes Project: AFR MAF=0.024; EUR MAF=0.042). Accordingly, we tested the phenotypic spectrum of rs201123820 together with the two SNPs (i.e., rs12461856 and rs9291211) that were identified in the European-ancestry and in the trans-ancestry meta-analyses, respectively. Twenty-six associations in our phenome-wide scan survived multiple testing correction accounting for the number of phenotypes and variants tested (FDR q<0.05; Figure 3; Supplementary Table 6). Among these, rs9291211 was associated with 22 phenotypes, which included dietary habits (e.g., UK Biobank Field ID: 6179 “*Mineral and other dietary supplements* [*None of the above*]”, p=1.68×10^−8^), anthropometric traits (e.g., UK Biobank Field ID: 1687 “*Comparative body size at age 10*”, p=7.58×10^−8^), behavioral traits (e.g., UK Biobank Field ID: 20127 “*Neuroticism score*”, p=3.12×10^−6^), physical outcomes (e.g., UK Biobank Field ID: 6152 “*Hay fever, allergic rhinitis or eczema*”, p=3.49×10^−5^), reproductive function (UK Biobank Field ID: 3581 *Age at menopause* [*last menstrual period*], p=8.40×10^−5^), and cognitive tests (e.g., UK Biobank Field ID: 404 “*Reaction time* [*Duration to first press of snap-button in each round*]”, p= 8.82×10^−5^). Rs201123820 was associated with several physical conditions: postprocedural musculoskeletal disorders (UK Biobank Field ID: 41202 “*Diagnoses – main ICD10* [*M96*]”, p=4.88×10^−7^), other disorders of the musculoskeletal system and connective tissue (UK Biobank Field ID: 41270 “*Diagnoses - ICD10* [*M13_MUSCULOSKELEOTH*]”, p=1.25×10^−5^), postpartum care and examination (UK Biobank Field ID: 41202 “*Diagnoses - main ICD10* [*Z39*]”, p=3.74×10^−5^), and auto-refraction measurements for eye prescription (UK Biobank Field ID: 5159 “*3mm asymmetry index (right)*”, p=6.54×10^−5^). No phenotypic associations with rs12461856 survived multiple testing correction.

**Figure 3:**
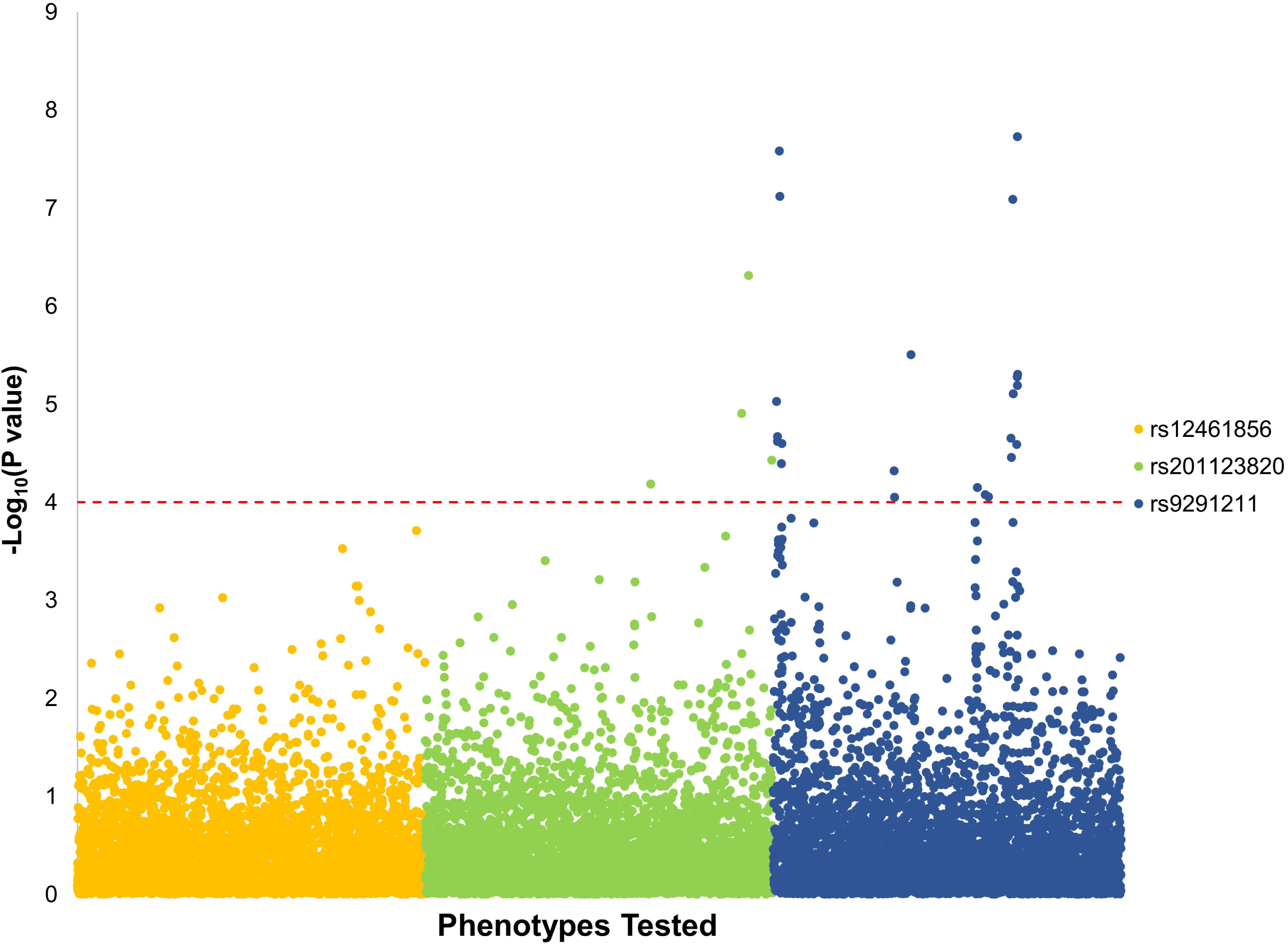
Manhattan plot of the phenome-wide scan conducted in the UK Biobank with respect to rs12461856, rs201123820, and rs9291211.

### Polygenic Risk Score Analysis

We also used PRS to compare the three opioid-related phenotypes – dependence (with exposed controls – OD_exposed_ – and with unexposed controls – OD_unexposed_) and exposure in non-dependent individuals (OE_controls_). Individuals exposed to opioids would be expected to have greater propensity to risk-taking behaviors than unexposed subjects^34^. Accordingly, we derived a PRS from the large-scale GWAS (N=466,571) conducted by the Social Science Genetic Association Consortium (SSGAC) on risk tolerance, which was defined as the tendency, preparedness, or willingness to take risks in general^31^. The PRS analysis was conducted on European-ancestry subjects only due to the well-known lack of large-scale GWAS in other ancestry groups^35^. The risk-tolerance PRS was positively associated with OD when contrasted with unexposed controls (OD_unexposed_: N_effective_=4,728, PT=1, z=3.94, p=8.1×10^−5^, FDR q=0.003), whereas OD contrasted with exposed controls displayed only a trend (p<0.1; OD_exposed:_ N_effective_=3,038, PT=1, z=1.93, p=0.054, FDR q=0.13). OE (OE_control_: N_effective_=5,376, PT=0.05, z=3.57, p=3.6×10^−5^, FDR q=0.003; Figure 4A) was also significant for the risk-tolerance PRS.

**Figure 4:**
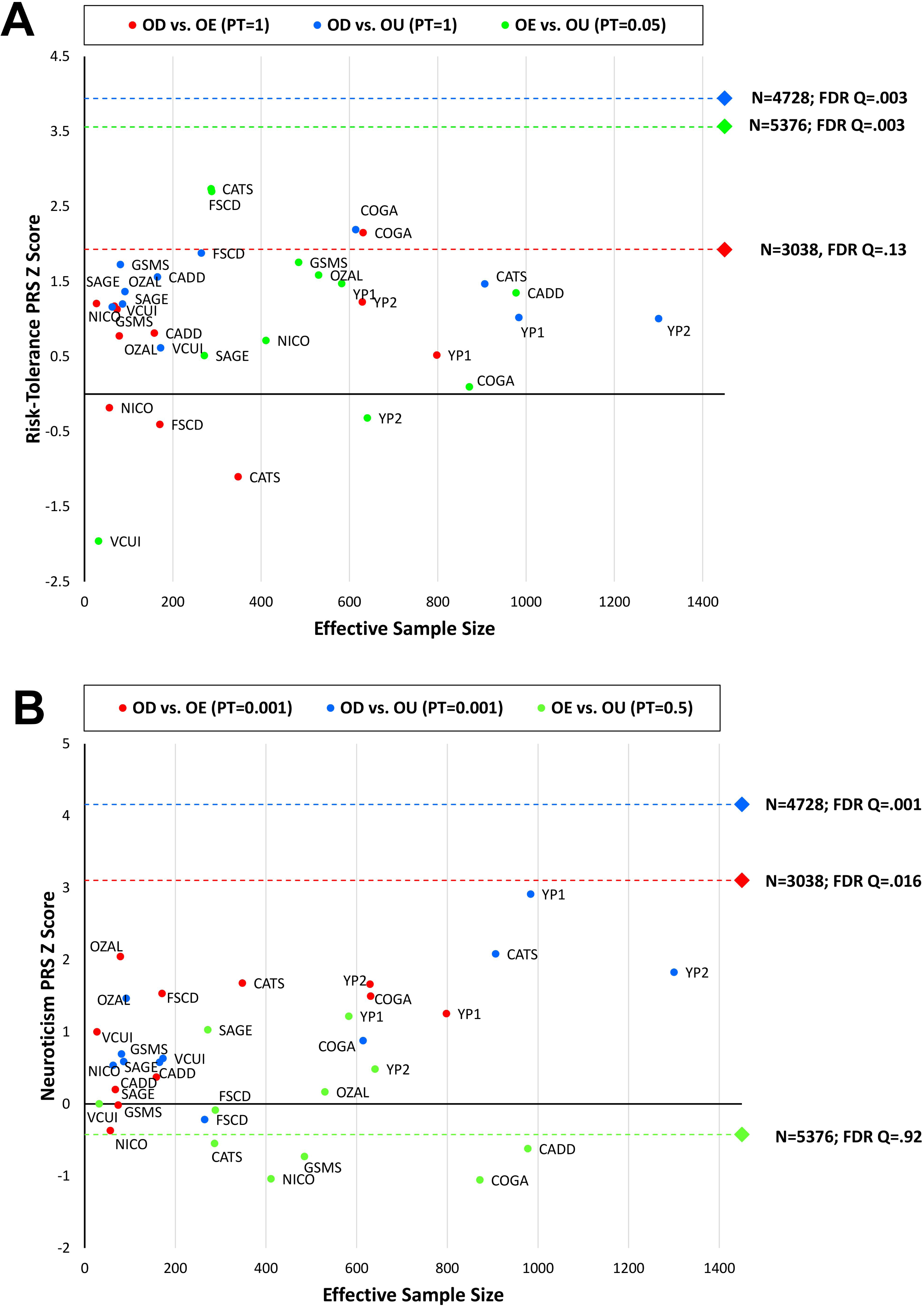
Relationship between PRS z scores (**A**: risk tolerance; **B**: neuroticism) and effective sample size across the opioid-related phenotypes tested. Each circle represents an individual cohort; the diamond represents the results from the meta-analysis with respect to the phenotypes tested.

We also tested PRS derived from a large-scale GWAS of neuroticism (N=390,278)^32^. This behavioral trait represents a tendency to negative affect and was previously observed to be genetically correlated with several psychiatric disorders, including SUDs^15, 36^, major depression^37^, and posttraumatic stress disorder^38^. Consistent with our expectation of genetic liability to negative affect being related to dependence but not exposure alone, the neuroticism PRS was associated with dependence compared with unexposed (OD_unexposed_: N_effective_=4,728, PT=0.001, z=4.16, p=3.2×10^−5^, FDR q=5.76×10^−4^) and dependence compared with exposed controls (OD_exposed_: N_effective_=3,038, PT=0.001, z=3.1, p=0.002, FDR q=0.016; Figure 4B), but not with exposed vs unexposed controls (OE_controls_: N_effective_=5,376, PT=0.5, z=-0.42, p=0.671, FDR q=0.919).

## DISCUSSION

We investigated the genetic architecture of opioid-related traits in informative cohorts. Our comparison of opioid dependence (OD) compared with unexposed (OD_unexposed_) and exposed (OD_exposed_) controls, as well as of opioid exposure among controls (OE_controls_) provides new insights into opioid addiction. We identified GWS loci and genes for OD_unexposed_ and OE_controls_, and found these variants to be associated with health relevant traits in the UK Biobank. Most critically, our PRS analyses highlighted distinctions between exposed and unexposed controls, as well as the progression from exposure to dependence. Despite shared biology, exposure and dependence to addictive substances are very different behaviors – one involves seeking or being exposed to a first use (and thereby cannot be related to the specific effect of the drug), while the other necessitates repeated use leading to the development of dependence. Use and dependence are behaviors with different relationships to other genetically influenced traits, as has been shown for alcohol use vs. alcohol dependence^39, 40^. The inclusion of unexposed individuals in the control group could introduce a bias toward to the null, to the extent there is a relatively large proportion of such controls who would have developed OD if they were exposed in the analyses. To our knowledge, no previous study investigated the specific genetic differences between OD and OE; our current findings provide the first insights based on genome-wide data into the molecular mechanisms by which OE and OD differ. The lack of sufficiently large numbers of OD cases and OE controls is a fundamental barrier to facilitating our understanding of the biological underpinnings of this serious public health epidemic, as it limited the power of what we would regard as the most informative comparison.

With respect to the single-variant associations observed, the strongest bioinformatics support from other studies was observed for rs9291211, identified in the European-ancestry GWAS meta-analysis of the OE_controls_ phenotype. Although this variant reached only a suggestive GWS threshold (p=7.2×10^−8^), we observed strong evidence of its regulatory effect on the brain-specific transcriptomic profile of *SLC30A9* and *BEND4*. The *BEND4* locus was also significantly confirmed by gene-based association analysis (p=9.9×10^−6^). The function of this gene is unclear, but previous GWAS identified several variants at this locus (including rs9291211) that were associated with depression^41^, alcohol consumption^33^, autism spectrum disorder^41^, neuroticism^32^, height^42^, and helping behavior^34^. *SLC30A9* encodes a zinc transporter involved in intracellular zinc homeostasis, which also plays a role in transcriptional activation of Wnt-responsive genes^43^. A rare 3 bp deletion in this gene (NM_006345.3, c.1047_1049delGCA, Ala350del) causes the recently discovered Birk-Landau-Perez syndrome, a combination of intellectual disability, muscle weakness, oculomotor apraxia, and nephropathy in early childhood^43^. In previous GWAS, variants located in *SLC30A9* gene (rs9291211 is located in *BEND4* but also affects *SLC30A9* gene expression) were associated with neuroticism^42^ and depression^44^. The phenome-wide scan of rs9291211 in the UK Biobank showed an effect of this SNP on a wide range of complex traits, some with an easy-to-conceptualize relationship to OD such as alcohol consumption, neuroticism, depression, and anxious feelings. However, the strongest results were observed with respect to dietary habits: rs9291211*A was positively associated with reduced OE risk in the PGC-SUD cohorts and with increased propensity to use dietary supplements, such as vitamin and mineral supplements in the UK Biobank. A recent GWAS identified several loci associated with dietary habits and indicated a causal relationship between educational attainment and healthy eating^45^. With respect to rs9291211, we also observed a nominally significant association with traits related to educational attainment (e.g., UK Biobank Field ID: 6138 *Qualification* [*College or University degree*], p=0.033). Accordingly, we hypothesize that rs9291211 could be involved in the individual variability to consume chemicals ranging from dietary supplements to opioids, independent from educational attainment. Beyond the intake of dietary supplements, rs9291211 was associated with several other phenotypic traits. Although some of them may be linked to OE and related traits, further studies will be needed to confirm the possible mechanisms explaining the convergence of these phenotypic associations.

In the trans-ancestry GWAS meta-analysis of the OE_controls_ phenotype comparison, we identified GWS loci in the single-variant and the gene-based analyses. No associations were observed for rs12461856 and further studies will be needed to confirm the validity of this finding. Conversely, *SDCCAG8* identified in the gene-based analysis (but not related to any individual GWS variants) was shown in studies available in the GWAS catalog^46^ to have 49 single-variant associations with educational attainment^47^, blood-related parameters^48^, risk-taking behaviors^31^, anthropometric traits^42^, kidney function^49^, and schizophrenia^50^. The previous associations with behavioral traits support *SDCCAG8* as potentially associated with behaviors that, in turn, associated with increased risk of OE.

With respect to the OD_unexposed_ phenotype comparison, we identified rs201123820 in the African-ancestry meta-analysis. This is a non-coding deletion located 2 kb upstream of *LOC101928144*, an uncharacterized long intergenic non-protein coding RNA. The gene-based association analysis identified a GWS locus in the same region, *C18orf32*, a gene involved in the activation of the NF-kappaB and MAPK signaling pathways, which play a key role in immune and inflammatory responses^51^. The phenome-wide analysis in the UK Biobank, despite its being in a predominantly European cohort, showed a significant association of rs201123820 with physical conditions, particularly musculoskeletal disorders. This is particularly interesting as opioids are commonly prescribed for pain management in musculoskeletal disorders^52^. Early use of opioids for musculoskeletal disorders is associated with prolonged work disability^52^, which may be related to the consequences of opioid abuse and/or the severity of the underlying disorder that required treatment. While these associations merit replication, this result highlights how human genetic research cannot only uncover behaviors related to OE and OD, but help to clarify the links between opioid use and abuse and the pain management protocols currently used clinically.

The available genome-wide data also permitted us to compare the opioid-related phenotypes with respect to shared genetic risk of relevant behavioral traits. Although there is variability in the effective sample size and therefore statistical power of the phenotypes tested (ODunexposed Neffective=4,728; OEcontrols: Neffective=5,376; ODexposed N_effective_=3,038), the PRS results showed an interesting pattern. The risk-tolerance PRS was positively associated with all three phenotypes with the strength of association mostly related to the effective sample size of the target sample. The association between OE_controls_ and genetic liability to risk-taking highlights the importance of accounting for the genetic factors related to the individual differences in exposure when examining those contribution to dependence. Future studies in larger samples could determine whether associations between risk-taking and OD_exposed_, which trended towards significance, are attenuated when compared with OD_unexposed_. Such a finding would support the hypothesis that the inclusion of exposed controls can “fine-tune” our ability to separate loci related to generalized risk-taking from those specific to repeated use that lead to opioid dependence. The neuroticism PRS showed positive associations with OD_unexposed_ and OD_exposed_ phenotype comparisons. Although it was non-significant, we observed a negative association of neuroticism PRS with the OE_controls_ phenotype. This suggestive negative relationship parallels the rs9291211 result where the allele A was associated with reduced OE in the PGC GWAS meta-analysis and increased neuroticism score in the UK Biobank. Genetic liability to neuroticism may thus overlap with genetic liability to OD but not to OE. In aggregate, the PRS findings show a continuum of association between opioid involvement and genetic liability to risk-taking but distinctions between exposure and dependence when considering genetic liability to negative affect. If confirmed, these findings would have major implications in the design of future SUD genetic research.

The present findings suggest that the genetics of opioid exposure, OE, differs in important ways from opioid dependence, OD, so that combining these groups can confound analyses. Fully understanding the genetic differences between OE and OD will require large, targeted collections of control samples that are carefully characterized as to types of exposure (medical and non-medical) to enable analysis of OE controls compared to OD cases.

Although several putative single-variant, gene-based, and PRS associations were identified based on the different OD and OE phenotypes, the sample size of the current investigation is small. The sample size was limited by the low prevalence of OD in the general population and by the societal stigma associated with it, which could decrease reporting. For these reasons, biobanks are likely to be less informative for these traits than for more socially acceptable substances, such as alcohol. Studies specifically targeting SUDs and assessing opioid-related behaviors are the only available samples that are informative for OD and OE GWAS, and to date there are few such large-scale studies available. This will limit the application of genetic instruments to clinical practice on SUDs. Another important limitation is the phenotypic heterogeneity within the opioid exposure sample, which included individual exposed to opioids via licit use (i.e., medical prescriptions) and illicit use. There may be important differences between these two subgroups (e.g., risk-taking may be more strongly associated with illicit exposure). However, several of the cohorts investigated lacked this information, and, due to the limited sample size, we were not able to make this comparison. In addition, this may have resulted in heterogeneity in the OU controls (i.e., those who reported not using opioids illicitly were unassessed for medical exposure). Finally, while the phenome-wide investigation in the UK Biobank provides encouraging support for the plausibility of our findings, it may reflect complex pleiotropic effects of these variants on multiple traits, including an unmeasured third variable. Replication of these association signals with opioid dependence and exposure phenotypes will be required.

In conclusion, we provide a comprehensive genome-wide investigation of opioid-related traits, highlighting different molecular mechanisms that could underlie exposure and dependence. These findings draw attention to challenges associated with the use of unexposed controls in genetic association studies for OD and potentially for other SUDs (where exposure is not widespread, as is the case for alcohol, or more recently marijuana). This information should be used to guide the next generation of human genetic studies of opioid-related behaviors.

## ACKNOWLEDGMENTS

The Psychiatric Genomics Consortium Substance Use Disorders Working Group receives support from the National Institute on Drug Abuse and the National Institute of Mental Health via U01 MH109532 and U01 MH109528. We gratefully acknowledge prior support from the National Institute on Alcohol Abuse and Alcoholism. Statistical analyses for the PGC were carried out on the Genetic Cluster Computer (http://www.geneticcluster.org) hosted by SURFsara and financially supported by the Netherlands Scientific Organization (NWO 480-05-003) along with a supplement from the Dutch Brain Foundation and the VU University Amsterdam.

AA acknowledges DA032573; ACH acknowledges support from NIH grants AA07535, AA07729, AA13320, AA13321, and AA11998; AEA acknowledges support from AA011408 and AA017828; AMG acknowledges support from U10 AA08401; BPR was supported by AA011408, AA017828 and AA022537; BTW acknowledges support from AA011408, AA017828 and AA022537; CJH acknowledges DA032555, DA035804, DA011015, and DA042755; DBH acknowledges support from R01DA036583; EJC acknowledges support from DA023026, DA011301, and DA024413; EOJ acknowledges support from R01 DA044014; HM acknowledges support from DA025109, DA024413, DA016977; JG acknowledges support from DA12690, DA047527; JKH acknowledges support from DA011015; KSK acknowledges support from AA011408, AA017828 and AA022537; LD is supported by an Australian National Health and Medical Research Council (NHMRC) Principal Research Fellowship; LJB acknowledges support from R01DA036583; LMH acknowledges support from AA011408 and AA017828; LMH acknowledges support from AA011408 and AA017828; MCS acknowledges support from DA035804; PAFM acknowledges funding support from NIH grants: DA012854 and R25DA027995; RAG acknowledges support from AA017444; REP is supported by NIH K01 grant MH113848; RP acknowledges support from DA12690 and DA047527; SAB acknowledges support from AA011408, AA017828, AA022537 and AA022717; SMH acknowledges support from R21AA024888 and K08DA032680; TBB acknowledges support from MH100549; TLW acknowledges support from R01 DA021905 and R01 DA035804; WEC acknowledges support from R01HD093651, R01DA036523, and P30DA023026, R01MH117559.

**Alcohol Dependence in African Americans (ADAA)** study was funded by NIH grant R01 AA017444. Funding support for the **Comorbidity and Trauma Study (CATS)** (dbGAP accession number: phs000277.v1.p1) was provided by the National Institute on Drug Abuse (R01 DA17305); GWAS genotyping services at the CIDR at The Johns Hopkins University were supported by the National Institutes of Health (contract N01-HG-65403). The data collection and analysis of the **Center on Antisocial Drug Dependence (CADD)** was supported by the following grants: DA011015, DA012845, DA021913, DA021905, DA032555, and DA035804. **The Collaborative Study on the Genetics of Alcoholism (COGA)** is supported by NIH Grant U10AA008401 from the National Institute on Alcohol Abuse and Alcoholism (NIAAA) and the National Institute on Drug Abuse (NIDA). Funding support for this GWAS genotyping, which was performed at the Johns Hopkins University Center for Inherited Disease Research, was provided by the National Institute on Alcohol Abuse and Alcoholism, the NIH GEI (U01HG004438), and the NIH contract “High throughput genotyping for studying the genetic contributions to human disease” (HHSN268200782096C). COGA Principal Investigators: B. Porjesz, V. Hesselbrock, H. Edenberg, L. Bierut, includes eleven different centers: University of Connecticut (V. Hesselbrock); Indiana University (H.J. Edenberg, J. Nurnberger Jr., T. Foroud); University of Iowa (S. Kuperman, J. Kramer); SUNY Downstate (B. Porjesz); Washington University in St. Louis (L. Bierut, J. Rice, K. Bucholz, A. Agrawal); University of California at San Diego (M. Schuckit); Rutgers University (J. Tischfield, A. Brooks); Department of Biomedical and Health Informatics, The Children’s Hospital of Philadelphia; Department of Genetics, Perelman School of Medicine, University of Pennsylvania, Philadelphia PA (L. Almasy), Virginia Commonwealth University (D. Dick), Icahn School of Medicine at Mount Sinai (A. Goate), and Howard University (R. Taylor). Other COGA collaborators include: L. Bauer (University of Connecticut); J. McClintick, L. Wetherill, X. Xuei, Y. Liu, D. Lai, S. O’Connor, M. Plawecki, S. Lourens (Indiana University); G. Chan (University of Iowa; University of Connecticut); J. Meyers, D. Chorlian, C. Kamarajan, A. Pandey, J. Zhang (SUNY Downstate); J.-C. Wang, M. Kapoor, S. Bertelsen (Icahn School of Medicine at Mount Sinai); A. Anokhin, V. McCutcheon, S. Saccone (Washington University); J. Salvatore, F. Aliev, B. Cho (Virginia Commonwealth University); and Mark Kos (University of Texas Rio Grande Valley). A. Parsian are the NIAAA Staff Collaborators. M. Reilly was an NIAAA staff collaborator. We continue to be inspired by our memories of Henri Begleiter and Theodore Reich, founding PI and Co-PI of COGA, and also owe a debt of gratitude to other past organizers of COGA, including Ting-Kai Li, currently a consultant with COGA, P. Michael Conneally, Raymond Crowe, and Wendy Reich, for their critical contributions. The authors thank Kim Doheny and Elizabeth Pugh from CIDR and Justin Paschall from the NCBI dbGaP staff for valuable assistance with genotyping and quality control in developing the dataset available at dbGaP (phs000125.v1.p1; also: phs000763.v1.p1; phs000976.v1.p1). Support for the **Study of Addiction: Genetics and Environment (SAGE)** was provided through the NIH Genes, Environment and Health Initiative [GEI; U01 HG004422; dbGaP study accession phs000092.v1.p1]. SAGE is one of the genome-wide association studies funded as part of the Gene Environment Association Studies (GENEVA) under GEI. Assistance with phenotype harmonization and genotype cleaning, as well as with general study coordination, was provided by the GENEVA Coordinating Center [U01 HG004446].

Assistance with data cleaning was provided by the National Center for Biotechnology Information. Support for collection of datasets and samples was provided by the Collaborative Study on the Genetics of Alcoholism [COGA; U10 AA008401], the **Collaborative Genetic Study of Nicotine Dependence** [COGEND; P01 CA089392; see also phs000404.v1.p1], and the **Family Study of Cocaine Dependence** [FSCD; R01 DA013423, R01 DA019963]. Funding support for genotyping, which was performed at the Johns Hopkins University Center for Inherited Disease Research (CIDR), was provided by the NIH GEI [U01HG004438], the National Institute on Alcohol Abuse and Alcoholism, the National Institute on Drug Abuse, and the NIH contract “High throughput genotyping for studying the genetic contributions to human disease” [HHSN268200782096C]. The **Gene–Environment Development Initiative: Great Smoky Mountains Study** (phs000852.v1.p1) was supported by the National Institute on Drug Abuse (U01DA024413, R01DA11301), the National Institute of Mental Health (R01MH063970, R01MH063671, R01MH048085, K01MH093731 and K23MH080230), NARSAD, and the William T. Grant Foundation. We are grateful to all the GSMS and CCC study participants who contributed to this work. The following grants supported data collection and analysis of CADD: DA011015, DA012845, DA021913, DA021905, DA032555, and DA035804. **Gene-Environment-Development Initiative -GEDI – Virginia Commonwealth University (VTSABD;** dbGAP in progress) was supported by the National Institute on Drug Abuse (U01DA024413, R01DA025109), the National Institute of Mental Health (R01MH045268, R01MH055557, R01MH068521), and the Virginia Tobacco Settlement Foundation grant 8520012. We are grateful to all the VTSABD-YAFU-TSA study participants who contributed to this work. **Yale-Penn** (phs000425.v1.p1; phs000952.v1.p1) was supported by National Institutes of Health Grants RC2 DA028909, R01 DA12690, R01 DA12849, R01 DA18432, R01 AA11330, and R01 AA017535 and the Veterans Affairs Connecticut and Philadelphia Veterans Affairs Mental Illness Research, Educational, and Clinical Centers. **Australian Alcohol and Nicotine studies (OZALC;** phs000181.v1.p1) were supported by National Institutes of Health Grants AA07535,AA07728, AA13320, AA13321, AA14041, AA11998, AA17688,DA012854, and DA019951; by Grants from the Australian National Health and Medical Research Council (241944, 339462, 389927,389875, 389891, 389892, 389938, 442915, 442981, 496739, 552485,and 552498); by Grants from the Australian Research Council(A7960034, A79906588, A79801419, DP0770096, DP0212016, and DP0343921); and by the 5th Framework Programme (FP-5) GenomEUtwin Project (QLG2-CT-2002-01254). Genome-wide association study genotyping at Center for Inherited Disease Research was supported by a Grant to the late Richard Todd, M.D., Ph.D., former Principal Investigator of Grant AA13320.

Substance Use Disorder Working Group of the Psychiatric Genomics Consortium: Raymond K. Walters, Renato Polimanti, Emma C. Johnson, Jeanette N. McClintick, Mark J. Adams, Amy E. Adkins, Fazil Aliev, Silviu-Alin Bacanu, Anthony Batzler, Sarah Bertelsen, Joanna M. Biernacka, Tim B. Bigdeli, Li-Shiun Chen, Toni-Kim Clarke, Yi-Ling Chou, Franziska Degenhardt, Anna R. Docherty, Alexis C. Edwards, Pierre Fontanillas, Jerome C. Foo, Louis Fox, Josef Frank, Ina Giegling, Scott Gordon, Laura M. Hack, Annette M. Hartmann, Sarah M. Hartz, Stefanie Heilmann-Heimbach, Stefan Herms, Colin Hodgkinson, Per Hoffmann, Jouke Jan Hottenga, Martin A. Kennedy, Mervi Alanne-Kinnunen, Bettina Konte, Jari Lahti, Marius Lahti-Pulkkinen, Dongbing Lai, Lannie Ligthart, Anu Loukola, Brion S. Maher, Hamdi Mbarek, Andrew M. McIntosh, Matthew B. McQueen, Jacquelyn L. Meyers, Yuri Milaneschi, Teemu Palviainen, John F. Pearson, Roseann E. Peterson, Samuli Ripatti, Euijung Ryu, Nancy L. Saccone, Jessica E. Salvatore, Sandra Sanchez-Roige, Melanie Schwandt, Richard Sherva, Fabian Streit, Jana Strohmaier, Nathaniel Thomas, Jen-Chyong Wang, Bradley T. Webb, Robbee Wedow, Leah Wetherill, Amanda G. Wills, 23andMe Research Team, Jason D. Boardman, Danfeng Chen, Doo-Sup Choi, William E. Copeland, Robert C. Culverhouse, Norbert Dahmen, Louisa Degenhardt, Benjamin W. Domingue, Sarah L. Elson, Mark A. Frye, Wolfgang Gäbel, Caroline Hayward, Marcus Ising, Margaret Keyes, Falk Kiefer, John Kramer, Samuel Kuperman, Susanne Lucae, Michael T. Lynskey, Wolfgang Maier, Karl Mann, Satu Männistö, Bertram Müller-Myhsok, Alison D. Murray, John I. Nurnberger, Aarno Palotie, Ulrich Preuss, Katri Räikkönen, Maureen D Reynolds, Monika Ridinger, Norbert Scherbaum, Marc A. Schuckit, Michael Soyka, Jens Treutlein, Stephanie Witt, Norbert Wodarz, Peter Zill, Daniel E. Adkins, Joseph M. Boden, Dorret I. Boomsma, Laura J. Bierut, Sandra A. Brown, Kathleen K. Bucholz, Sven Cichon, E. Jane Costello, Harriet de Wit, Nancy Diazgranados, Danielle M. Dick, Johan G. Eriksson, Lindsay A. Farrer, Tatiana M. Foroud, Nathan A. Gillespie, Alison M. Goate, David Goldman, Richard A. Grucza, Dana B. Hancock, Kathleen Mullan Harris, Andrew C. Heath, Victor Hesselbrock, John K. Hewitt, Christian J. Hopfer, John Horwood, William Iacono, Eric O. Johnson, Jaakko A. Kaprio, Victor M. Karpyak, Kenneth S. Kendler, Henry R. Kranzler, Kenneth Krauter, Paul Lichtenstein, Penelope A. Lind, Matt McGue, James MacKillop, Pamela A. F. Madden, Hermine H. Maes, Patrik Magnusson, Nicholas G. Martin, Sarah E. Medland, Grant W. Montgomery, Elliot C. Nelson, Markus M. Nöthen, Abraham A. Palmer, Nancy L. Pedersen, Brenda W. J. H. Penninx, Bernice Porjesz, John P. Rice, Marcella Rietschel, Brien P. Riley, Richard Rose, Dan Rujescu, Pei-Hong Shen, Judy Silberg, Michael C. Stallings, Ralph E. Tarter, Michael M. Vanyukov, Scott Vrieze, Tamara L. Wall, John B. Whitfield, Hongyu Zhao, Benjamin M. Neale, Joel Gelernter, Howard J. Edenberg & Arpana Agrawal

