## Supplemental Material for "Leveraging genome-wide data to investigate differences between opioid use *vs*. opioid dependence in 41,176 individuals from the Psychiatric Genomics Consortium"

**Supplementary Methods**

*Cohorts*

Alcohol Dependence in African Americans (ADAA) data were collected between 2009 and 2013 and consisted of cases recruited from treatment centers in St. Louis Missouri and controls screened for the absence of substance use disorders recruited from households selected from neighborhoods in proximity to neighborhoods of residence of case participants. Cases met criteria for DSM-IV opioid dependence. Controls were opioid-exposed but did not meet criteria for opioid dependence (DSM-IV).

Australian Alcohol and Nicotine Studies (OZAL) study recruited participants from twins and their relatives who had participated in questionnaire- and interview-based studies on alcohol and nicotine use and alcohol-related events or symptoms^1^. They were living in Australia and of predominantly European ancestry Opioid dependence was defined using DSM-IV criteria.

Center on Antisocial Drug Dependence (CADD) cohort includes unrelated participants aggregated from several studies described elsewhere^2-5^. This cohort was over-selected for adolescent behavioral disinhibition, with half of the participants ascertained specifically from high-risk populations (i.e. recruited through substance abuse treatment, special schools, or involvement with the criminal justice system; see supplement of 19 for additional criteria for clinical probands). Lifetime opioid dependence was assessed with the CIDI-SAM and defined as meeting opioid dependence at any wave for this longitudinal study.

Collaborative Study on the Genetics of Alcoholism (COGA) is a multi-site study of alcohol dependent probands and their family members. Alcohol dependent probands were recruited from inpatient and outpatient facilities. Community probands and their family members were also recruited from a variety of sources. Further details were described previously ^6^. All participants were assessed using the Semi-Structured Assessment for the Genetics of Alcoholism (SSAGA). Cases met criteria for a lifetime history of DSMIV opioid dependence. The exposed controls were defined as those who were exposed at least once in their lifetime to opioids but did not meet criteria for opioid dependence.

Comorbidity and Trauma Study (CATS) consisted of opioid dependent individuals aged 18 and older recruited from opioid substitution therapy clinics in the greater Sydney area and genetically unrelated individuals with little or no lifetime opioid misuse from neighborhoods in geographic proximity to these clinics. All subjects were of European-Australian descent. Further details were described previously^7^. All participants were assessed using the SSGA. Opioid dependence was defined using DSM-IV criteria. The exposed controls were defined as those who were exposed at least once in their lifetime to opioids but did not meet criteria for opioid dependence at the time of their last interview.

Gene-Environment-Development Initiative – the Great Smoky Mountains Study (GSMS) is prospective study funded by the National Institute on Drug Abuse (NIDA). The Participants were assessed via structured interviewing using the Young Adult Psychiatric Assessment and its early life extension (i.e., YAPA and CAPA), yielding diagnoses and symptom scales for a wide range of substance use disorders (SUDs). Opioid dependence was defined using DSM-IV criteria. Exposed controls were defined as those who were exposed to opioids at least once but did not meet criteria for opioid dependence.

Gene-Environment-Development Initiative – Virginia Commonwealth University (VCUI) study combined existing phenotypic and environmental data from the Virginia Twin Study of Adolescent Behavioral Development (VTSABD) study, a population-based multi-wave, cohort-sequential twin study of adolescent psychopathology and its risk factors, and two followup studies, the Young Adult Follow Up (YAFU) and the Transitions to Substance Abuse (TSA) study. Further details of the VCUI cohort were reported previously^8, 9^. Participants were assessed via structured interviewing using the Child Adult Psychiatric Assessment (CAPA), a Structured Clinical Interview for DSM-IV (SCID)-based assessment of psycho-pathology in young adult twins for YAFU and the Life Experiences Interview (LEI) for TSA, yielding diagnoses and symptom scales for a wide range of substance use disorders (SUDs). Opioid dependence was defined using DSM-IV criteria.

Study of Addiction: Genetics and Environment (SAGE), Collaborative Genetic Study of Nicotine Dependence (COGEND) and Family Study of Cocaine Dependence (FSCD) are selected from three large, complementary studies focused of the genetics of substance use disorders^10-12^. We analyze these subsets separately and remove overlap between cohorts. SAGE participants were assessed using the SSAGA. FSCD and COGEND participants were assessed using polydiagnostic instruments closely based on the SSAGA. Cases reported a lifetime history of DSM-IV opioid dependence. Genetically unrelated control subjects reported to be exposed to opioids at least once in their lifetime but had no significant opioid-dependence symptoms.

Yale-Penn (YP) study includes participants recruited in the eastern US, predominantly in Connecticut and Pennsylvania. They were administered the Semi-Structured Assessment for Drug Dependence and Alcoholism (SSADDA) to derive DSM-IV diagnoses of lifetime substance use disorders and other major psychiatric traits). The study received IRB approval from all participating institutions and written informed consent was obtained from all study participants. Additional information is available in the previous GWAS publications^13-17^. Opioid dependence was defined using DSM-IV criteria.

*Inflation and Deflation in the cohorts investigated*

Genomic inflation factor (λ_GC_) and LD score regression intercept were calculated to investigate the presence of the deviations of the genome-wide association results from the expected distribution. Inflation (λ_GC_>1.04) could be due to the effect of unaccounted population stratification and polygenicity^18^. To distinguish polygenicity from unaccounted population stratification, we calculated the LD score regression intercept in the datasets tested^19^. Additionally, we observed in some of the cohorts investigated a strong deflation (λ_GC_<0.9), which is due to the low number of cases and the small case-control ratio present in some of the cohorts. Accordingly, we excluded NICO, OZAL, SAGE, and YP2 (AFR) from the OD_unexposed_ meta-analysis due to the deflation observed.

**Supplementary Table 1**: Sample size of the opioid-informative cohorts investigated.

| **Cohort** | | **Ancestry** | **Cases** | **Exposed**  **Controls** | **Unexposed**  **Controls** |
| --- | --- | --- | --- | --- | --- |
| **case-control studies: logistic regression** | | | | | |
| COGEND Study of Addiction: Genetics and Environment (SAGE) | | EUR | 22 | 73 | 966 |
| Collaborative Study on the Genetics of Nicotine Dependence (NICO) | | EUR | 15 | 117 | 794 |
| Center on Antisocial Drug Dependence (CADD) | | EUR | 44 | 386 | 666 |
| Gene–Environment Development Initiative: Great Smoky Mountains Study (GSMS) | | EUR | 21 | 151 | 615 |
| Family Study of Cocaine Dependence (FSCD) | | AFR | 57 | 66 | 477 |
|  |  | EUR | 81 | 98 | 361 |
| Comorbidity and Trauma Study (CATS) | | EUR | 1025 | 95 | 291 |
| Alcohol Dependence in African Americans (ADAA) | | AFR | 241 | 177 | 1393 |
| **Family-based studies: logistic mixed model** | | | | | |
| Yale-Penn | Phase1 (YP1) | AFR | 626 | 580 | 1840 |
|  |  | EUR | 1030 | 295 | 426 |
|  | Phase2 (YP2) | AFR | 204 | 214 | 952 |
|  |  | EUR | 642 | 210 | 685 |
| Gene–Environment Development Initiative: Virginia Commonwealth University (VCUI) | | EUR | 48 | 10 | 2482 |
| Australian Alcohol and Nicotine Studies (OZAL) | | EUR | 24 | 173 | 12088 |
| Collaborative Study on the Genetics of Alcoholism (COGA) | | AFR | 103 | 260 | 2401 |
|  |  | EUR | 320 | 1268 | 6063 |
| **TOTAL** | | | **4,503** | **4,173** | **32,500** |

**Supplementary Table 2:** Number of variants tested in the ancestry-specific and trans-ancestry meta-analysis across the phenotypes investigated. The association analysis was conducted considering variants present in at least 80% of the cohorts investigated.

| **Meta-analysis** | **OD_exposed_** | **OD_unexposed_** | **OE_controls_** |
| --- | --- | --- | --- |
| **African-ancestry** | 8,956,178 | 8,896,928 | 9,065,342 |
| **European-ancestry** | 4,211,587 | 4,887,055 | 5,986,959 |
| **Trans-ancestry** | 4,000,397 | 4,010,688 | 5,122,696 |

**Supplementary Table 3**: Genomic inflation factors (λ_GC_) across the meta-analyses conducted with respect to different phenotypic definitions and ancestry groups. LD score regression intercept was calculated when λ_GC_ > 1.04 to distinguish polygenicity from population stratification.

| **Meta-analysis** | **OD vs. OE** | **OD vs. OU** | **OE vs. OU** |
| --- | --- | --- | --- |
| African-ancestry | 0.994 | 1.018 | 0.999 |
| European-ancestry | 1.033 | 1.055  (LDSC intercept=1.033) | 1.044  (LDSC intercept=1.020) |
| Trans-ancestry | 1.028 | 1.061  (LDSC intercept=1.029) | 1.031 |

**Supplementary Table 4:** Ancestry-specific results for each genome-wide significant variant identified.

| **Phenotype** | **Meta-analysis** | **rsID** | **Effect Allele** | **Other Allele** | **Effect Allele Frequency** | **Z score** | **P value** |
| --- | --- | --- | --- | --- | --- | --- | --- |
| OD_unexposed_ | AFR | rs201123820 | T | TAAACAAAAACA | 0.0193 | 5.547 | 2.90E-08 |
|  | EUR |  |  |  | 0.0402 | -0.903 | 0.3665 |
|  | TRANS |  |  |  | 0.0318 | 2.472 | 0.01345 |
| OE_controls_ | TRANS | rs12461856 | A | G | 0.8402 | -5.606 | 2.07E-08 |
|  | EUR |  |  |  | 0.8436 | -4.833 | 1.35E-06 |
|  | AFR |  |  |  | 0.833 | -2.866 | 0.004163 |
|  | EUR | rs9291211 | A | G | 0.7817 | -5.387 | 7.16E-08 |
|  | AFR |  |  |  | 0.331 | 0.829 | 0.407 |
|  | TRANS |  |  |  | 0.6356 | -3.956 | 7.61E-05 |

**Supplementary Table 5:** directions and heterogeneity estimates of the three GWS variants among the cohorts meta-analyzed.

| **rsID** | **Effect Allele** | **Other**  **Allele** | **Z score** | **P value** | **Direction** | **Heterogeneity** | | | |
| --- | --- | --- | --- | --- | --- | --- | --- | --- | --- |
|  |  |  |  |  |  | **I^2^** | **Χ^2^** | **Df** | **P value** |
| rs201123820 | T | TAAACAAAAACA | 5.547 | 2.90E-08 | +++++ | 0 | 3.534 | 4 | 0.472 |
| rs9291211 | A | G | -5.387 | 7.16E-08 | -------?--? | 0 | 3.74 | 8 | 0.880 |
| rs12461856 | A | G | -5.606 | 2.07E-08 | ------??-?----?- | 0 | 9.744 | 11 | 0.554 |

**Supplementary Table 6:** Results in the present study related to variants identified in prior opioid dependence GWAS. We report the results of the variants that survived the quality control criteria applied to the different meta-analysis (e.g., information present in at least 80% of the cohorts meta-analyzed; imputation info score > 0.8; minor allele frequency > 0.01).

| Phenotype | Ancestry | rsID (Gene, Reference) | Effect Allele | Other Allele | Effect Allele Frequency | Z score | P value |
| --- | --- | --- | --- | --- | --- | --- | --- |
| OE_controls_ | TRANS | rs10494334 (Intergenic, ref.10) | A | G | 0.1035 | 0.439 | 0.6605 |
| OE_controls_ | EUR |  | A | G | 0.0677 | 0.336 | 0.7367 |
| OD_exposed_ | AFR |  | A | G | 0.1851 | -0.23 | 0.818 |
| OE_controls_ | TRANS | rs12442183 (*RGMA*, ref.9) | T | C | 0.3793 | -1.709 | 0.08753 |
| OE_controls_ | EUR |  | T | C | 0.408 | -1.819 | 0.06886 |
| OD_exposed_ | AFR |  | T | C | 0.3202 | 1.043 | 0.2968 |
| OE_controls_ | TRANS | rs1436175 (*CNIH3*, ref.7) | A | G | 0.4638 | 0.049 | 0.9609 |
| OE_controls_ | EUR |  | A | G | 0.4092 | 0.085 | 0.9325 |
| OD_exposed_ | AFR |  | A | G | 0.5901 | 0.269 | 0.7881 |
| OD_exposed_ | TRANS |  | A | G | 0.4861 | -1.827 | 0.0677 |
| OD_exposed_ | EUR |  | A | G | 0.4264 | -2.496 | 0.01256 |
| OD_unexposed_ | TRANS |  | A | G | 0.4771 | -0.786 | 0.4319 |
| OD_unexposed_ | EUR |  | A | G | 0.4054 | -0.869 | 0.3847 |
| OE_controls_ | TRANS | rs62103177 (*KCNG2*, ref.8) | A | G | 0.1188 | 0.988 | 0.323 |
| OE_controls_ | EUR |  | A | G | 0.1436 | 0.5 | 0.6173 |
| OD_exposed_ | AFR |  | A | G | 0.0725 | -1.625 | 0.1043 |
| OD_exposed_ | TRANS |  | A | G | 0.1159 | -2.09 | 0.03664 |
| OD_exposed_ | EUR |  | A | G | 0.1456 | -1.368 | 0.1712 |
| OD_unexposed_ | TRANS |  | A | G | 0.1165 | -1.666 | 0.09574 |
| OD_unexposed_ | EUR |  | A | G | 0.1458 | -1.557 | 0.1195 |

**Supplementary Table 7**: full summary association data of the phenome-wide scan conducted in the UK biobank. [xlsx file attached].

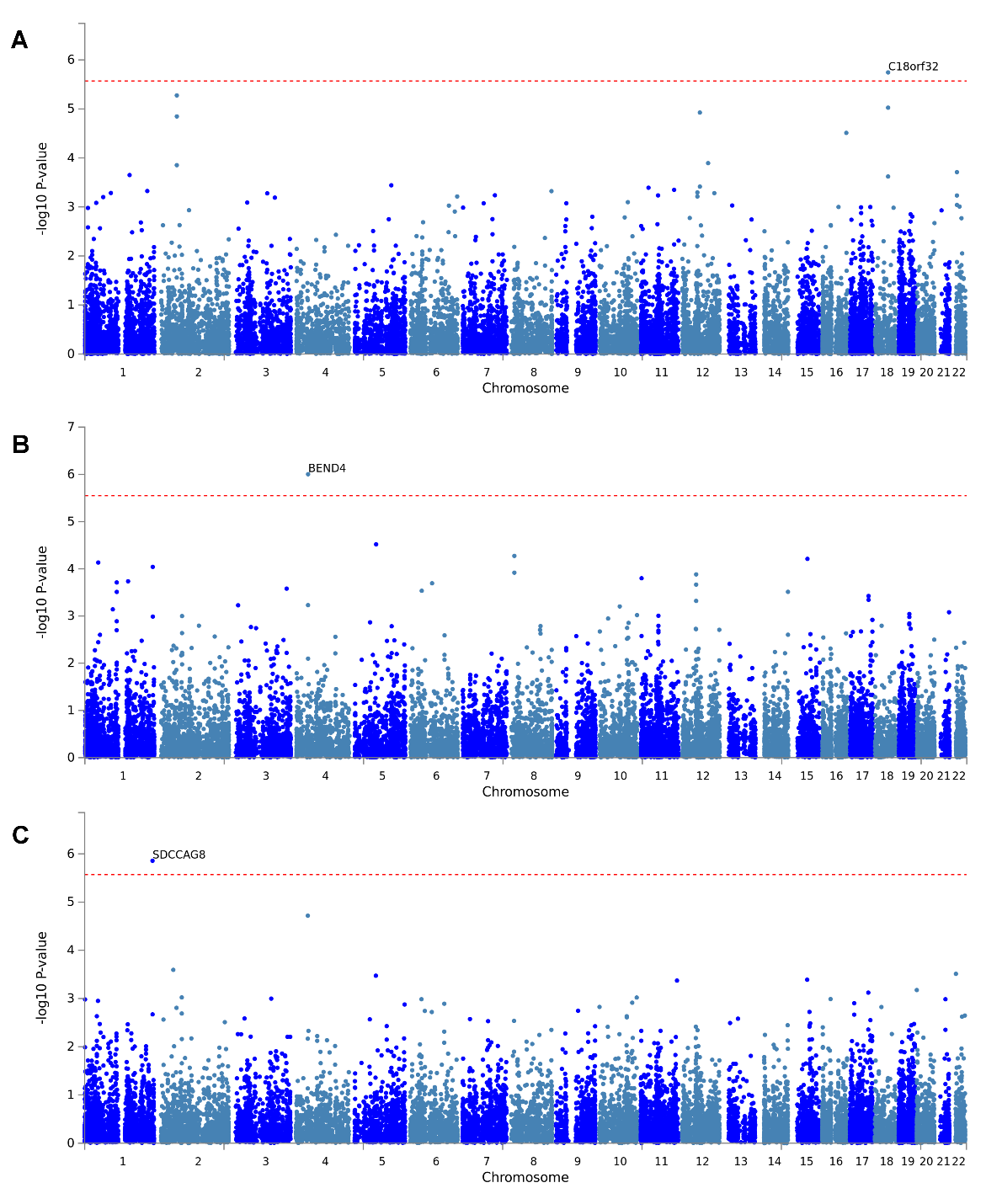

**Supplementary Figure 1**: Manhattan plots from the gene-based GWAS meta-analysis of OD_unexposed_ phenotype in African-ancestry individuals (**A**); OE_controls_ phenotypes in European-ancestry individuals (**B**) and in the trans-ancestry meta-analysis (**C**).

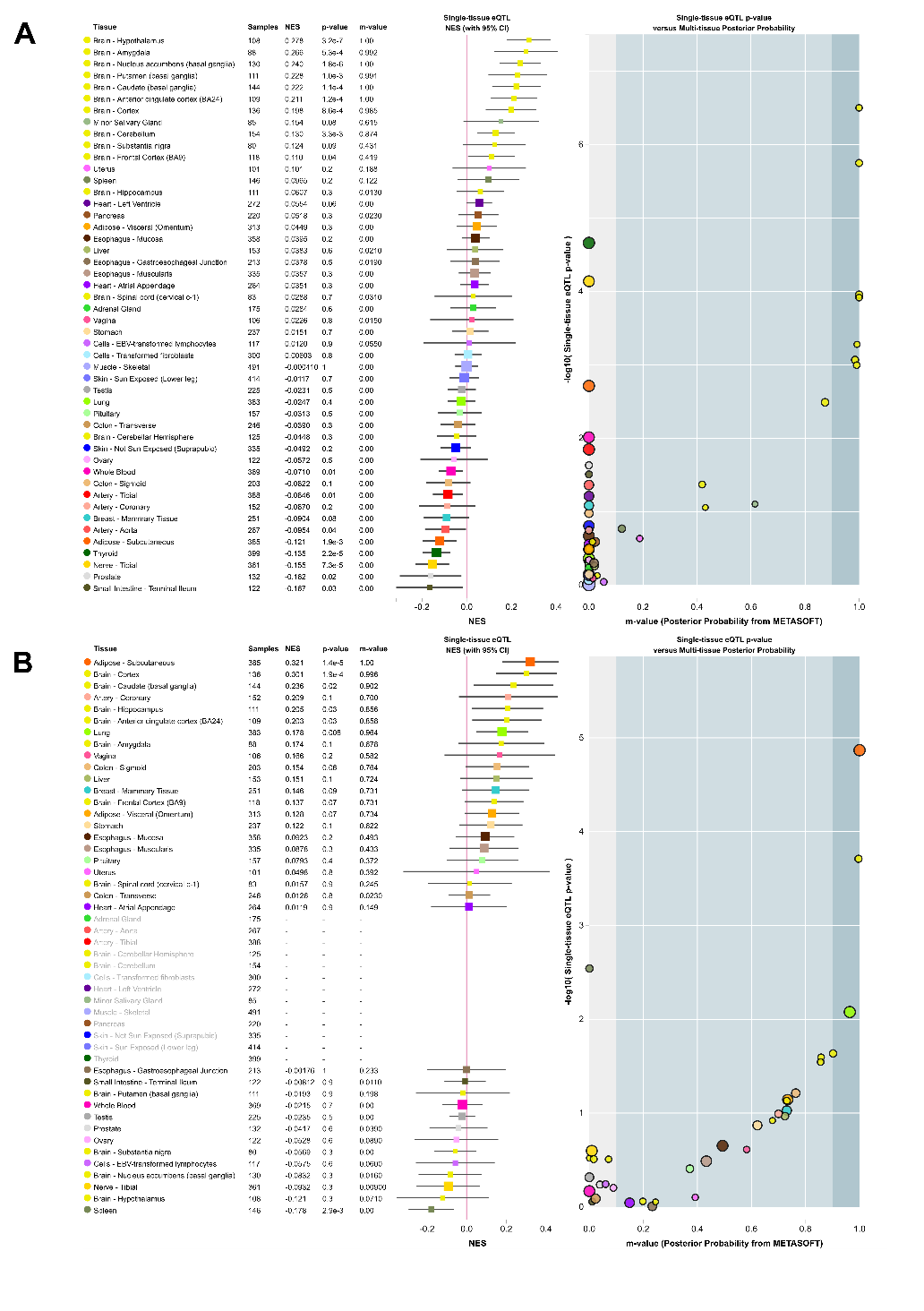

**Supplementary Figure 2**: Multi-tissue eQTL results of rs9291211 with respect to *SLC30A9* (**A**) and *BEND4* (**B**) transcriptomic profiles.
